# N-Terminal proteomics reveals distinct protein degradation patterns in different types of human atherosclerotic plaques

**DOI:** 10.1101/2024.05.14.594251

**Authors:** Lasse G. Lorentzen, Karin Yeung, Nikolaj Eldrup, Jonas P. Eiberg, Michael J. Davies

## Abstract

**BACKGROUND:** Destabilization and rupture of atherosclerotic plaques is a major cause of acute atherosclerotic cardiovascular events, including heart attack, ischemic stroke and peripheral arterial disease. Plaque destabilization is associated with extracellular matrix (ECM) modification and remodelling involving protease activity. Enzymatic cleavage generates protein fragments with new ‘ends’ (N-termini). We hypothesized that plaques susceptible to rupture would contain elevated levels of fragmented proteins with new N-termini. Identification of active proteases and their target proteins might allow categorization of plaque stability.

**METHODS:** Plaques from 21 patients who underwent carotid surgery due to symptomatic carotid artery stenosis were examined in an observational/cross-sectional study. The plaques were solubilized, digested, enriched for N-terminal fragments and analyzed by liquid chromatography-mass spectrometry.

**RESULTS:** The above methodology detected 35349 peptides, with 19543 being N-terminal species; 6561 were subsequently identified and quantified. Multidimensional scaling analysis and hierarchical clustering indicate the presence of three distinct clusters, which correlate with gross macroscopic plaque morphology (soft, mixed, and hard), ultrasound classification (echolucent/echogenic) and presence of hemorrhage/ulceration. Major differences were identified in the complement of peptide fragments, consistent with alternative turnover and degradation pathways dependent on plaque type. Identified peptides include signal and pro-peptides from ECM synthesis/turnover, and many from protein fragmentation. Sequence analysis indicates the targeted proteins (including ECM species) and the proteases (including meprins, cathepsins, matrix metalloproteinases, elastase, kallikreins) involved in fragment generation.

**CONCLUSIONS:** This study provides a large dataset of peptide fragments and proteases involved in plaque stability, mechanistic insights into remodelling, and possible biomarkers for improved atherosclerosis risk profiling.

**GRAPHICAL ABSTRACT:** 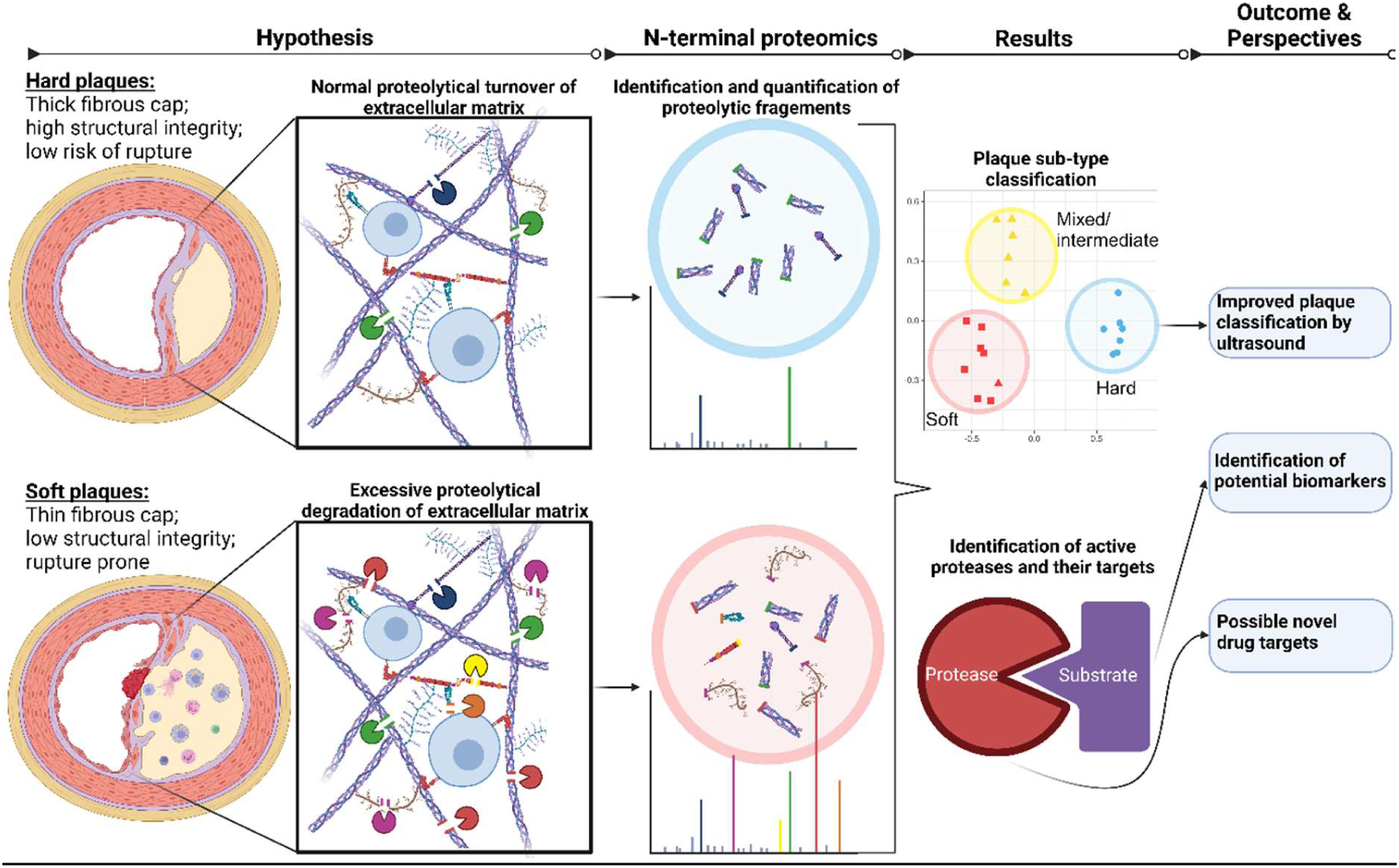

---

Atherosclerosis is characterized by the thickening and hardening of arterial walls due to accumulation of cholesterol, fat, and other substances in the vessel wall. Atherosclerosis and CVD are intricately interconnected, as the build-up of atherosclerotic plaques can induce ischaemia and embolization in organs such as the heart and brain. Consequently, atherosclerosis gives rise to adverse cardiovascular complications, including symptomatic ischaemic heart disease, cerebrovascular disease, and peripheral arterial disease ^1^. The stability of atherosclerotic plaques is critical factor in the severity of the condition, as progression can lead to unstable plaques which rupture and trigger acute and severe cardiovascular events ^2,3^.

Understanding the mechanisms underlying plaque progression and stability is crucial for developing effective therapeutic interventions ^2,3^. Plaque instability and the subsequent risk of cardiovascular events are closely linked to the integrity of the fibrous cap, a protective layer consisting of endothelial and smooth muscle cells, collagen, elastin, and other ECM components covering the lipid-rich core of atherosclerotic plaques. This cap serves as a mechanical barrier, preventing the exposure of the thrombogenic lipid-rich and necrotic core to the bloodstream.

Whilst protease activity is critical for ECM synthesis, as this requires the removal of signaling and pro-peptides to allow correct ECM assembly, unplanned or excessive ECM degradation or removal can compromise the structural integrity of the fibrous cap ^4,5^ resulting in destabilization, erosion, ulceration and embolization ^3^. Enzyme-mediated proteolysis (e.g. collagen and elastin) is reported to play a critical role, with this mediated by matrix metalloproteinases (MMPs), the disintegrin and metalloproteinase family (ADAMs), cathepsins and others ^4–6^.

Inflammatory cells, such as macrophages, neutrophils and T lymphocytes, release proteases via ongoing inflammatory responses within the plaque ^2–5,7^. Most proteases are secreted in inactive pro-forms, with activation triggered by pro-inflammatory cytokines, chemokines and other proteases within the plaque microenvironment ^2,3,8^. Oxidative stress, which is enhanced in atherosclerotic plaques ^9,10^, can promote protein damage and degradation via direct reactions, but also indirectly by altering protease levels and activity. Thus oxidants and their products can stimulate the expression, processing and secretion of proteases ^11,12^. Oxidants can also activate MMPs and inhibit their endogenous inhibitors (e.g. tissue inhibitors of matrix metalloproteinases, TIMPs) ^13–15^. Understanding the interplay between proteolysis and fibrous cap destabilization is therefore essential for developing therapeutic strategies to stabilize vulnerable plaques and prevent cardiovascular events.

The present study utilized cutting-edge mass spectrometry-based degradomic techniques to identify protein fragments and responsible proteases, to unravel the mechanisms underlying proteolysis in atherosclerotic plaques. This proteomics approach allows the identification of protease substrates and their cleavage sites within stable and unstable plaques.

## METHODS

An extended description of the methods employed in provided in the Supplementary Data.

### Data Availability Statement

The authors declare that all data are available within the article, in the Supplementary content or, for the MS data, available via ProteomeXchange with identifier PXD052236. See also the Major Resources Table in the Supplemental Materials.

### Inclusion criteria and sample size

Carotid plaques were collected from 21 consecutive patients undergoing carotid endarterectomy (CEA) due to symptomatic carotid stenosis (SCS) defined as recent ischaemic stroke (cerebral emboli) or recent retinal ischaemia (retinal artery emboli). The total proteome of the plaques from these subjects has been reported previously.^16^ Patients were diagnosed, initially medically treated and subsequently referred for vascular surgery. All patients were subsequently referred to the same referral center of vascular surgery for CEA. Demographic and other data on the subjects is provided in Supplementary Table 1.

### Ultrasound (US) imaging

US imaging of carotid plaques was done 1-3 days preoperatively by one MD and researcher with substantial experience in carotid US (KY, co-author). Data were obtained as 10-second video clips (cine loops) in sweeping axial and longitudinal planes to capture the whole carotid plaque. Based on the US acquisitions, plaques were categorized using a modified Gray-Weale classification into one of five catagories ^17^: uniformly echolucent (type 1), predominantly echolucent with < 50% echogenic areas (type 2), predominantly echogenic plaque with < 50% echolucent areas, uniformly echogenic (type 4) and plaques that could not be classified because of heavy calcification (type 5) ^17–19^. The US morphology classification was determined by the same researcher performing the US examinations (KY, co-author), who was blinded to the perioperative macroscopic classification and proteomic data.

### Tissue collection and processing

The sample collection and processing workflow was as described previously ^16^. Plaques were removed *in toto,* and then proteins were extracted using the SPEED protocol ^20^. Protein extracts were reduced and alkylated followed by clean-up using paramagnetic beads ^21^. The proteins were then digested with TrypN and the resulting peptides desalted using C18 cartridges, then subjected to strong cation exchange separation on self-packed stagetips ^22,23^. The eluted fractions were then analysed by LC-MS/MS.

### LC-MS/MS acquisition

Peptides were separated by reversed-phase C18 column chromatography using a Dionex Ultimate RSLCnano system wth gradient elution, then analysed on a timsTOF Pro mass spectrometer (Bruker) Data was acquired using the DDA-PASEF mode.

### Data analysis

MSFragger was used to search against a human proteome database digested *in silico* N-terminally to Arg and Lys residues, while allowing for non-specific cleavage at the peptide N-terminal. Modification of the N-terminus by acetylation, carbamylation or carboxymethylation was included as a variable modification together with Met oxidation and pyro-Glu formation. Search results were filtered using a FDR of 1% at precursor and peptide level. Peptide signals were quantified with match-between-runs enabled within FragPipe. Analysis of differential abundance of N-terminal peptides between plaques was carried out using a robust linear model approach. Proteins were considered as differentially expressed if the Benjamini–Hochberg adjusted p-value ^24^ was < 0.01 and the log2 fold change was above 1 or below -1.

Over-representation analysis of GO (*molecular function*) terms was carried out using the clusterProfiler package with parent proteins as input and a Benjamini–Hochberg adjusted p-value of < 0.05 as cut-off. The TopFind database was used to map N-terminal peptides to known cleavage and translation initiation sites.

### Ethical considerations

The study was conducted according to the Helsinki declaration, and data and biological materials were collected after patients’ written, verbal, and informed consent. Study approval was obtained from The Danish National Committee on Health Research Ethics (journal number H-20002776).

## RESULTS

Twenty-one atherosclerotic plaques were analyzed using a TrypN-based degradomics approach (Figure 1A). A total of 35349 peptides were identified, of which 19543 were N-terminal peptides. After filtering to remove peptides with missed cleavages and high numbers of missing values (i.e. detected in a limited numbers of samples), 5815 unique peptide sequences (6561 peptidoforms in total, when post-translational modifications are included) were selected for further investigation (Figure 1B,C; a full list of peptides is given in Supplementary Tables 2 and 3) and compared against annotations in the TopFind database ^25,26^. This mapping revealed distinct categories of peptides, and different proteins from which these originate within the plaques. A number of the peptides (621) mapped to canonical start sites, indicating identification of proteins at their expected initiation sites. Twenty-one additional peptides were associated with alternative splicing sites, consistent with splice variants. Most (4388) were identified as neo-N-terminal peptides, indicating that they arise from proteolytic cleavage events that give new N-terminal sequences. Of these, 312 were identified as pro-peptides and 244 as signal peptides, indicating the presence of protein precursors that undergo proteolytic processing to yield mature forms; these data are consistent with ongoing ECM and intracellular protein synthesis. Of the identified N-terminal peptides, 229 were identified as arising from proteins (mainly immunoglobulins) not present in the TopFind database, and thus could not be mapped (Figure 1D).

**Figure 1:**
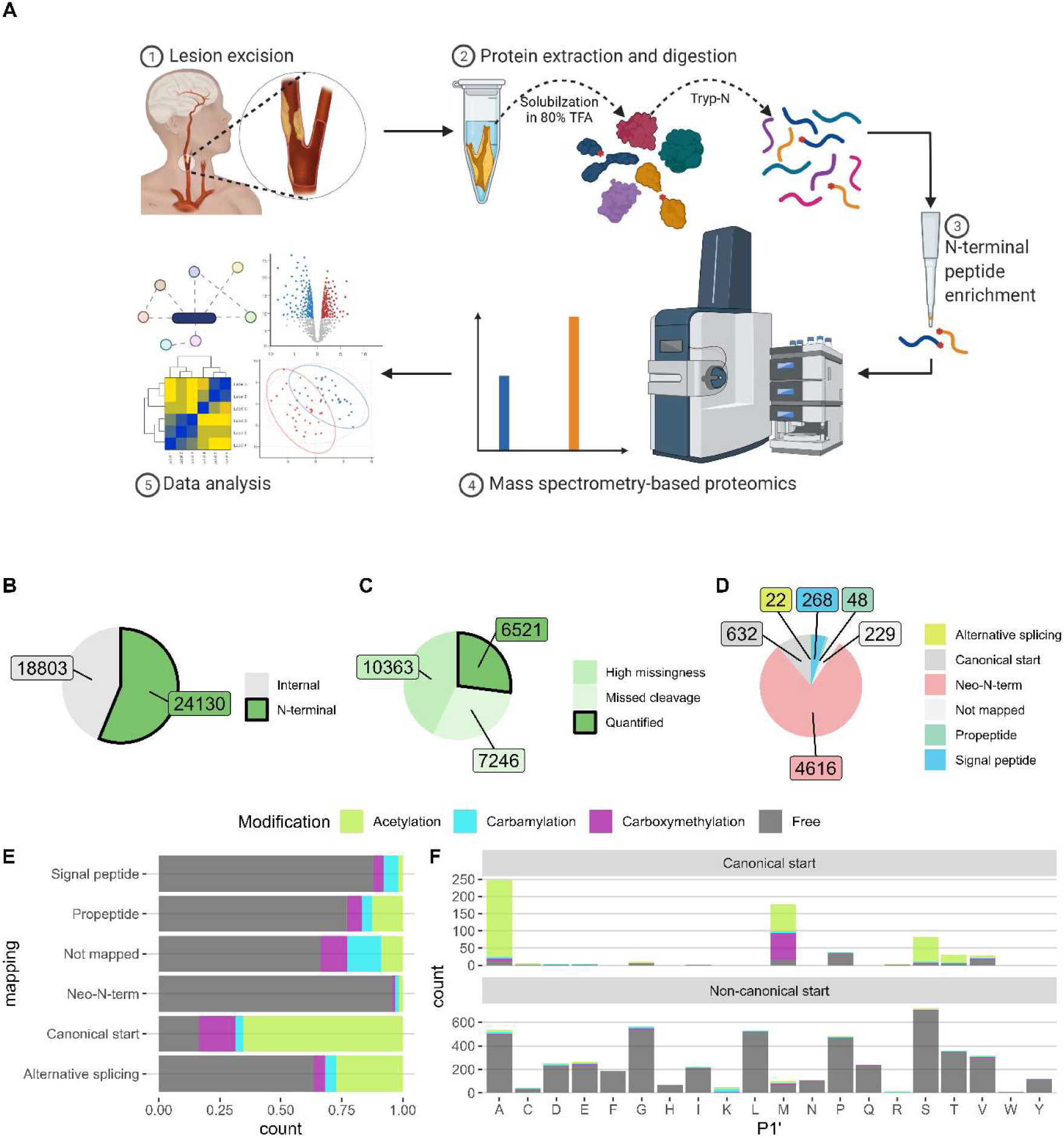
N-Terminal proteomics workflow and data summary. **(A)** Schematic representation of the N-terminal proteomics workflow: (1) Plaques obtained during carotid endarterectomy were subjected (2) to protein extraction and a clean-up process using magnetic bead capture and digestion using the protease Tryp-N. (3) The N-terminal peptides were subsequently enriched through the capture of internal peptides on strong-cation exchange material. (4) These peptides were then identified and quantified via LC-MS/MS and, (5) analyzed for abundance, protein origin, positional mapping, sites of cleavage and potential differences between plaques. **(B)** Pie chart indicating the distribution of identified N-terminal and internal peptides of the species detected. **(C)** Pie chart illustrating the number of quantified N-terminal peptides compared to those excluded on the basis of missed cleavages or high levels of missing data. **(D)** Pie chart showcasing the distribution of the 6,521 quantified N-terminal peptides, categorized by their mapping to canonical start sites, neo-N-terminal species, alternative splicing, pro-peptides, signal peptides and species that could not be mapped. **(E)** Chart providing the percentage of native (free amino group) and modified N-terminal peptides for each type of mapping. **(F)** Bar plot displaying the count of N-terminal peptides, categorized by the amino acid at the P1’ position (i.e. the N-terminal amino acid), with these color-coded based on the presence of a free N-terminal amino group or the presence of modifications at the initial amino acid. The upper panel represents N-terminal peptides mapping to canonical start sites (and indicating the high extent of modifications present at such start sites), while the lower panel presents the data for the species identified as non-canonical N-terminal peptides. Data are from 21 carotid plaques from 21 individuals.

Acetylation is a common co- or post-translational modification associated with N-terminal amines. We identified 1395 such species, of which 516 were quantified. Most (1283) mapped to canonical protein initiation or alternative splicing sites. An open search showed that carbamylation and carboxymethylation were also prevalent N-terminal modifications, which were included in a closed database search. We identified 916 carboxymethylated and 885 carbamylated N-terminal peptides, of which 183 and 150 could be quantified, respectively. Acetylation was primarily observed for canonical N-terminal peptides with Ala or Met at P1’ (with P1’ indicating the new N-terminus, P2’ the second amino acid, etc), and Ser and Thr to a lesser extent (top part of Figure 1F). Carboxymethylation was primarily detected at N-terminal Met and to a lesser extent at Ala. Carbamylation was less prevalent, and detected on all canonical N-terminal residues except Pro (Figure 1F). In contrast, multiple amino acids were detected as the N-terminal residues in peptides with non-canonical start sites (lower part of Figure 1F) and these had low levels of acetylation, carbamylation or carboxymethylation.

Quantitative label-free quantification values from the 5815 identified and unique N-terminal peptides were used for consensus clustering (k-means algorithm ^27^). This revealed that the 21 plaques clustered optimally into three distinct groups (Figure 2A-C). Multidimensional scaling (MDS) demonstrated clear segregation of Clusters 1 and 2, with Cluster 3 between these, but closer to Cluster 2 than 1 (Figure 2B). This analysis was compared to hierarchical clustering and other biological data for these samples, with the latter analyses carried out independently and in a blinded manner. The proteomic clusters correlated strongly with plaque morphology (categorized as hard, mixed, soft; determined through gross perioperative macroscopic examination of the plaques) and the presence of ulceration and thrombi (Figure 2C). Strong and significant overlaps were also observed with US grayscale classifications obtained prior to surgery (Figure 2C). The excellent agreement between proteomic cluster assignment and plaque morphology suggests that the identified clusters capture underlying differences in plaque characteristics. Furthermore, these data indicate that specific and distinct clusters of N-terminal peptides are present in plaques of different types.

**Figure 2:**
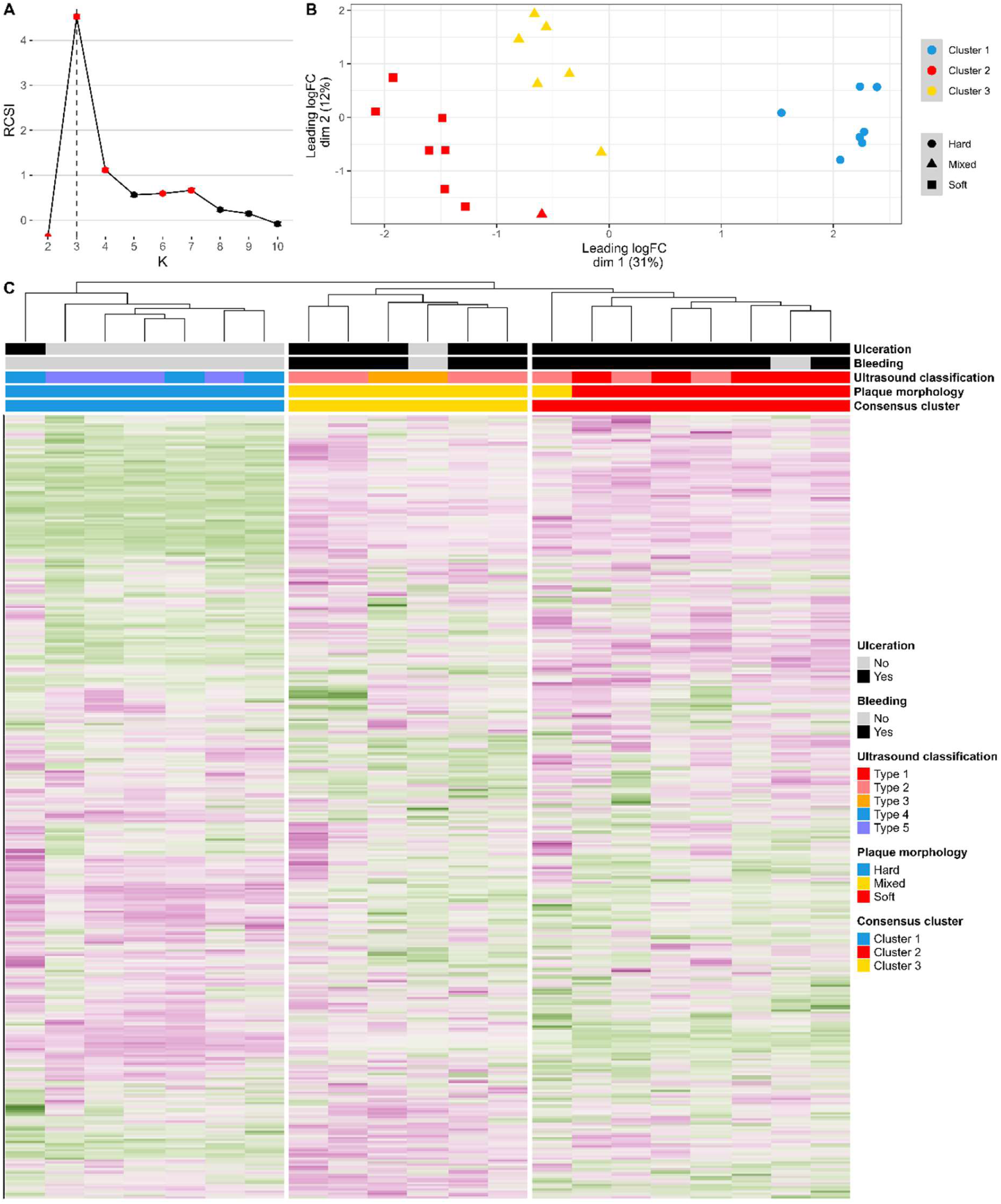
Cluster analysis and profiling of N-terminal peptides. **(A)** The relative cluster stability index for a range of cluster numbers (K) from 2 to 10, obtained through Monte Carlo reference-based consensus clustering. Clusters with significantly different stability scores from the null distribution (p < 0.01) are highlighted in red. The vertical line indicates the optimal cluster number determined by the M3C algorithm. **(B)** Multidimensional scaling (MDS) plot representing the distances between abundance profiles of N-terminal peptides. Each sample is displayed on a two-dimensional scatterplot using the first two components of the MDS analysis. Each dot corresponds to an individual plaque, color-coded based on their cluster assignment for the K = 3 cluster (from (A). **(C)** Top section: consensus clustering of the plaques with regards to the presence of erosion, bleeding, ultrasound classification and gross plaque morphology. Bottom section: heat map providing a visualization of the scaled abundances of all quantified N-terminal peptides (rows) for each of the carotid plaques analyzed (columns) with the dendogram indicating Euclidian distances between plaque samples. These data (from 21 carotid plaques from 21 individuals) indicate that there are distinct N-terminal peptide clusters within the plaque samples, with these aligning with the consensus clustering based on the presence of erosion and bleeding, ultrasound classification and gross plaque morphology.

Linear modeling was used to further examine differences in the abundance of the N-terminal peptides between the clusters. This revealed that 1527 N-terminal peptides were significantly different in abundance across at least one of the three clusters (Figure 3A). Specifically, 284 N-terminal peptides displayed significantly increased abundance in Cluster 2, and 731 were significantly more abundant in Cluster 2 and 3. Similarly, 325 were significantly more abundant in Cluster 1, and 97 N-terminal peptides were more abundant in both Clusters 1 and 3 (Figure 3B,C). These data are consistent with the MDS data. The majority of the significantly-different N-terminal peptides mapped to neo-N-termini (Figure 3D). In addition, 216 canonical N-terminal peptides were found to be significantly different between all 3 clusters, with most of these peptides being acetylated forms.

**Figure 3:**
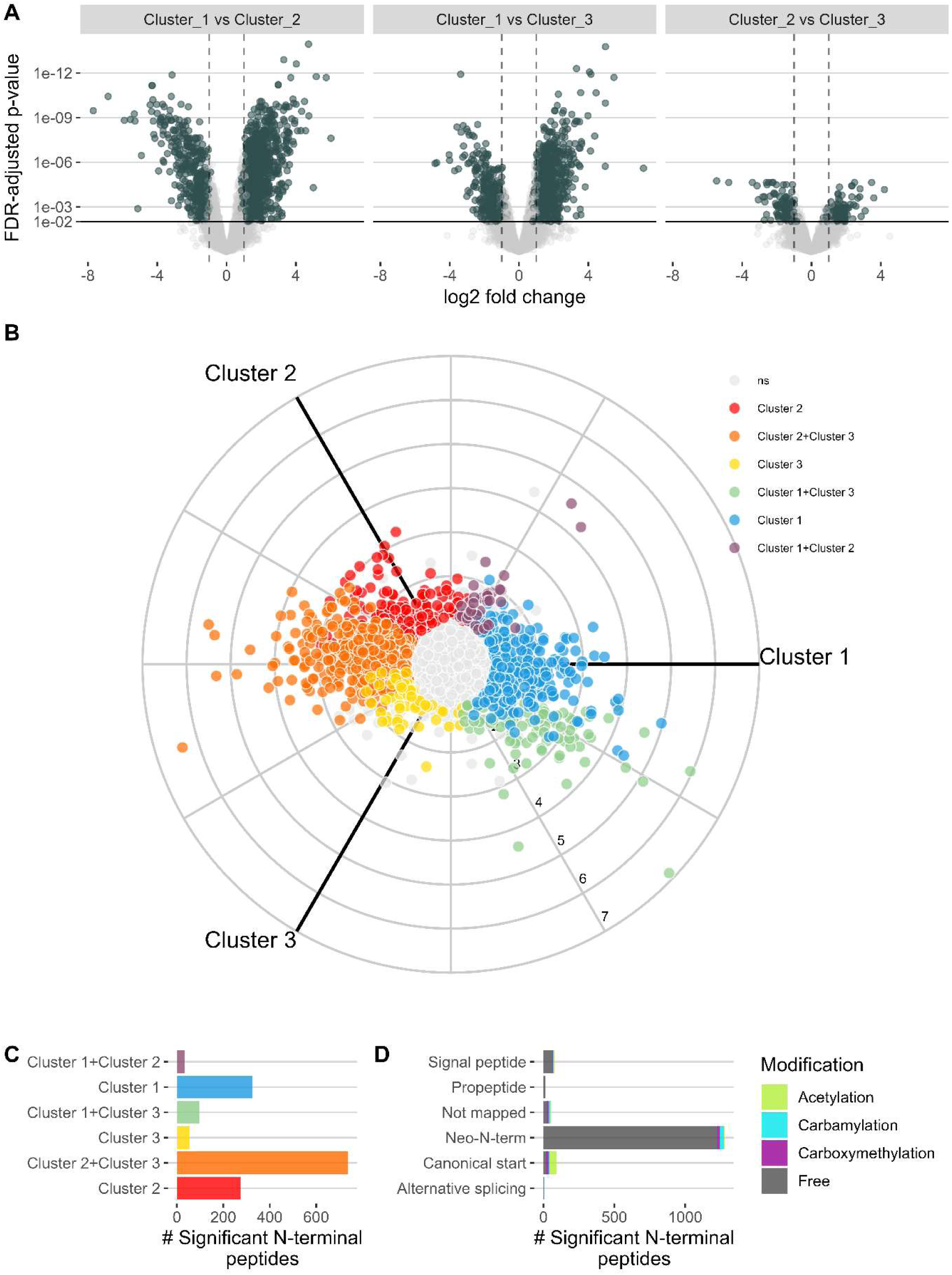
Comparative analysis of N-terminal peptides in different plaque types. **(A)** Volcano plot of the N-terminal peptides and their abundance compared between the 3 different clusters using robust linear modelling. Each data point corresponds to an N-terminal peptide, with the log2 fold change between compared samples on the x-axis and the false discovery rate (FDR)–adjusted P values (-log10 transformed) on the y-axis. The non-axial vertical lines indicate ±1 fold change, and the non-axial horizontal line denotes the significance cut-off at adjusted P value > 0.01, such that peptides indicated by dark grey symbols are significantly differentially expressed, and those in light grey are not. **(B)** Three-axis polar plot depicting the abundance of N-terminal peptides, color-coded to indicate significantly increased abundance in each cluster, and combinations of these clusters (i.e. plaque types). N-terminal peptides that did not display differential abundance in any of the three clusters (adjusted p value > 0.01) are depicted as grey dots in the centre of the plot. **(C)** Bar plot illustrating the count of N-terminal peptides that exhibit significantly increased abundance in each cluster, and combinations of the cluster sets. **(D)** Bar plot showing the number of N-terminal peptides with significantly different abundances in any of the three clusters, categorized by their mapping within the protein sequence, and color-coded based on their modifications. Data are from 21 carotid plaques from 21 individuals.

The analysis of differentially-abundant N-terminal peptides revealed distinct patterns of protein representation between Clusters 1 and 2. Among the identified proteins, 29 were represented by 10 or more N-terminal peptides that displayed significant differences in abundance between any of the three clusters (Figure 4A). These 29 proteins accounted for 689 N-terminal peptides that exhibited significant abundance variations between any of the 3 clusters, and >40% of the total number of significantly different N-terminal peptides (Figure 4B). These findings indicate that modest numbers of proteins are strongly associated with the observed differences in the numbers of neo-N-terminal peptides detected in each cluster, and between the different clusters. However, other proteins with fewer differentially-abundant N-terminal peptides may contribute to a more subtle or localized impact on the observed cluster-specific proteolytic profiles.

**Figure 4:**
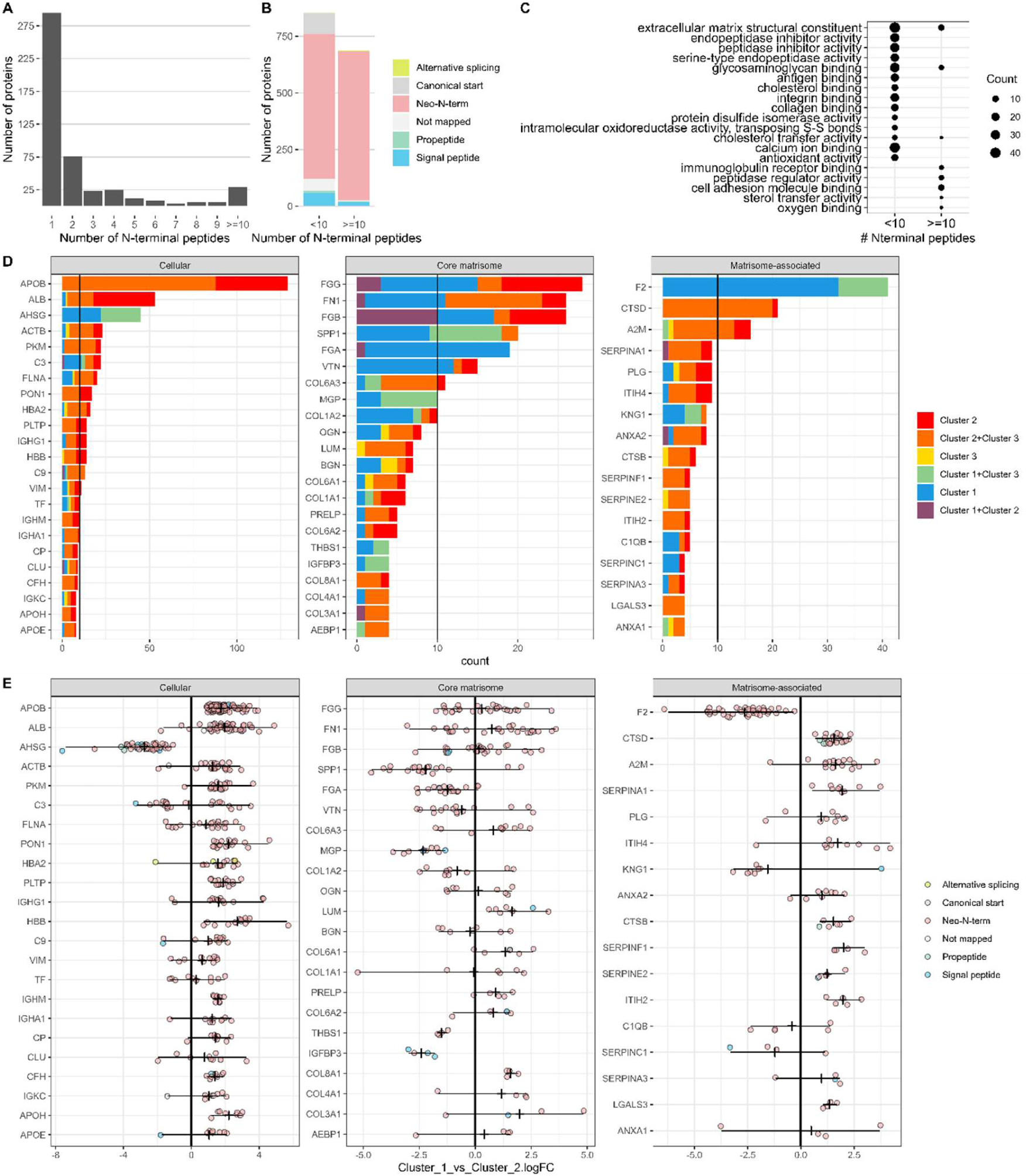
Protein and N-terminal peptide abundance across plaque types. **(A)** Bar plot displaying the number of proteins detected as giving rise to 1-9, or 10 and more, N-terminal peptides that are significantly upregulated in any of the three clusters. **(B)** Bar plot indicating the count of significant N-terminal peptides categorized by their origin (neo-N-terminal etc), from proteins giving rise to either < 10, or > 10, N-terminal peptides, color-coded to the origin of the peptides from the protein sequence. **(C)** Dot plot presenting Gene Ontology (GO) enrichment terms for proteins giving rise to < 10, and 10 or greater, N-terminal peptides. **(D)** Bar plot showing the number of N-terminal peptides with significantly increased abundance in any of the three clusters, categorized by the source proteins and color-coded to the cluster(s) from which they detected as being over abundant. Three panels display data for proteins categorized as cellular (left), core matrisome (middle), and matrisome-associated (right). For ease of display, only proteins yielding with the largest numbers of N-terminal peptides are included (cellular proteins > 7; core and matrisome-associated > 3). **(E)** Jitter plot indicating the relative change in N-terminal peptide abundances between Clusters 1 and 2, categorized by their source proteins. Only N-terminal peptides that exhibit significant differences (adjusted P value < 0.01) are presented. Points are color-coded according to the mapped location of the N-terminal peptide in the protein sequence. Data are from 21 carotid plaques from 21 individuals.

GO term enrichment analysis of the proteins yielding >10 significant N-terminal peptides, versus proteins with <10 such peptides, revealed both shared and distinct functional categories. ECM proteins, glycosaminoglycan-binding proteins and proteins involved in the regulation of proteolytic activity were strongly enriched, indicating that these are major targets (Figure 4C). The proteins that yielded the largest numbers of neo-N-terminal peptides are displayed in Figure 4D,E. These proteins are divided into separate classes (cellular, core matrisome and matrisome-associated) and color coded to the specific plaque clusters. Apolipoprotein B contributed the most neo-peptides in the cellular division, with this abundance being particularly associated with clusters 2 and 3 (soft and mixed).

In order to investigate the active proteases that generate the neo-N-terminal peptides, the topFIND database ^26^ was used to search for protease-substrate interactions. A total of 567 were matched to a known protease cleavage sites, with 159 differing significantly between the 3 clusters.

Among the proteases identified, several stand out. These include meprins 1B and 1A (MEP1B, MEP1A), cathepsins (B, D, S, and L isoforms; CATB, CATD, CATS, CATL1), matrix metalloproteases (MMP3, MMP14, MMP7, MMP11, MMP13, MMP12), chymase (CMA1) and neutrophil elastase (ELNE) (Figure 5A). Several of these have multiple substrates that are differentially abundant between the 3 clusters (Figure 5A), and are particularly abundant in clusters 2 and 3 (e.g. meprins and cathepsins, in soft and mixed plaques), whereas others (e.g. MMP14, MMP13 and MMP8) are primarily associated with cluster 1 (hard plaques). This indicates that proteases are active in all plaque sub-types, but that these and their substrates differ between the clusters (Figure 5B).

**Figure 5:**
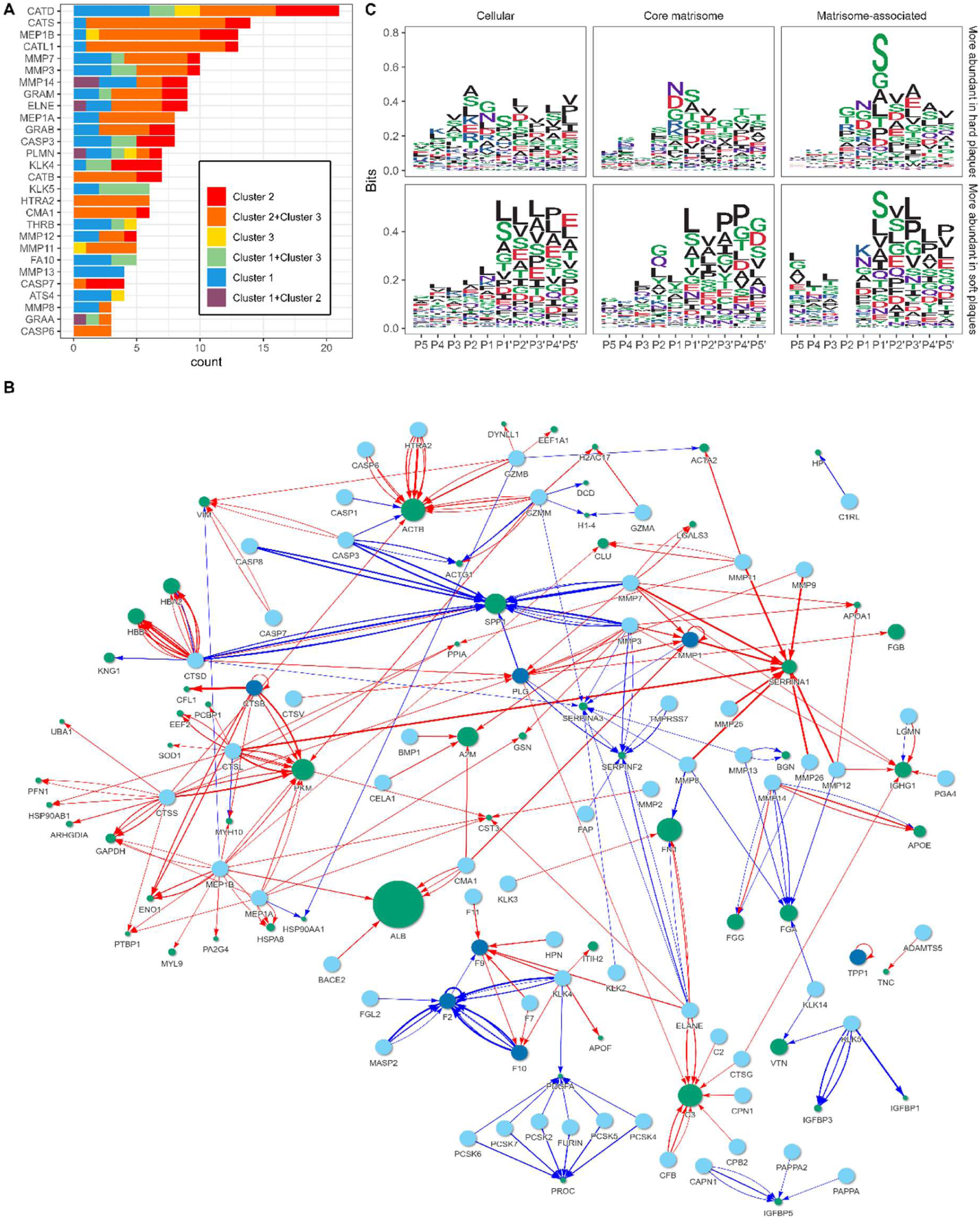
Proteases responsible for generation of detected neo-N-terminal peptides, identification of proteolytic sequence specificity, and substrate-protease interactions. **(A)** Bar plot displaying the count of neo-N-terminal peptides with significantly increased abundance in any of the three clusters, categorized by the cleaving proteases according to the TopFIND database. Bars are color-coded based on the cluster(s) where the neo-N-terminal peptides are more abundant. **(B)** Network plot illustrating protease-substrate interactions according to the TopFIND database. Light-blue nodes represent cleaving proteases, and green nodes represent substrates. Connecting lines represent a neo-N-terminal peptide from that protein where the cleavage site has previously been linked to the protease in question. Lines are color-coded according to the cluster in which the neo-N-terminal peptides are more abundant (red: more abundant in cluster 2, blue: more abundant in cluster 1), and the line width is proportional to the log2 fold change. The size of the substrate nodes represents the total number of neo-N-terminal peptides from that protein identified as differentially abundant between clusters 1 and 2. **(C)** Sequence logo plots for neo-N-terminal peptides identified as more abundant in cluster 1 (upper panels) or cluster 2 (lower panels). Separate panels are displayed for proteins assigned as cellular (left), core matrisome (middle), and matrisome-associated proteins (right). Data are from 21 carotid plaques from 21 individuals.

While the topFIND analysis provided valuable data, many of the total and neo-N-terminal peptides were not present in this database. As an alternative, sequence logo analysis was performed to uncover patterns in cleavage specificity. An enrichment of Leu (L, and to a lesser extent Pro, P) was observed at, or around, P1’ (the new N-terminus) for N-terminal peptides more abundant in clusters 2 or 3 (Figure 5C, lower part). In contrast, enrichment for Leu or Pro was not detected for the peptides in cluster 1 (Figure 5C, top part). This enrichment was present for all protein divisions (cellular, core matrisome and matrisome associated). In contrast, Ser was more enriched at P1’ for N-terminal peptides more abundant in Cluster 1. These differences are protease dependent, and consistent with different enzymes being involved in homeostasis and disease.

## DISCUSSION

In the current study, peptides generated from plaque proteins as a result of fragmentation (the plaque ‘degradome’) have been identified by N-terminal proteomics ^22,23^ using digestion with TrypN, and subsequent separation of endogenous peptides from TrypN-generated species uisng strong cation exchange resin. This approach enables precise mapping of biologically-generated N-termini resulting from protease cleavage, and identification of proteases and target proteins present within plaque sub-types. This approach has allowed identification of many thousands of peptides, of which 5815 were examined further. Sequence analysis unveiled groups of peptides with distinct origins. Most were neo-N-terminal peptides consistent with *in vivo* proteolysis of plaque proteins.

Cluster analysis based on label-free quantitation values, provided evidence for three distinct clusters with with cluster 3 falling between cluster 1 and 2, but closer to cluster 2. These clusters exhibited strong correlations with plaque morphology, as determined through macroscopic assessment and ultrasound classification. These therefore represent distinct plaque phenotypes recognizable by imaging techniques, underscoring the clinical relevance of the data.

Significant numbers of N-terminal peptides were differentially abundant between the clusters, yielding data on the proteolytic profiles of plaque phenotypes. Many identified peptides were strongly associated with protein differences between clusters 1 and 2, as evidenced by multiple, differentially-abundant peptides derived from them. GO term enrichment analysis indicated that ECM proteins, glycosaminoglycan-binding proteins, and proteins involved in regulating proteolytic activity were strongly enriched, highlighting their importance as targets.

Meprins, cathepsins, neutrophil elastase and some MMPs have been identified as important in generating peptides from proteins in cluster 2 (soft) plaques, consistent with an increased activity of these species in this phenotype (Figures 3 and 4). With few exceptions, proteases are excreted as inactive forms that require pro-peptide removal (or alteration) for activation ^28,29^; the processes that drive activation in plaques are however unclear (see below). In contrast to the data for cluster 2, MMPs 8, 13 and 14 appear to be more active in cluster 1 (hard) plaques. This proteolytic activity and resulting peptides, may arise from ECM synthesis and maintenance of a thick fibrous cap.

Some of the identified proteases have been previously linked with atherosclerosis. The gene *Mep1a*, which codes for meprin 1A (MEP1A), a member of the metzincin (metalloendopeptidase, astacin) family, is a susceptibility gene for atherosclerosis in apoE^-/-^ mice ^30^ but this has not been reported previously for humans ^31^. Neither MEP1A, nor its isoform MEP1B also detected here, have been reported previously as being elevated in human plaques. Meprin inhibition in apoE^-/-^ mice reduces plaque size ^32^ decrease cell apoptosis, and down-regulates oxidant production ^32^. MEP1B can influence immune cell activity ^31,33^, upregulate important pro-inflammatory factors including interleukin 1-beta (IL-1β), IL-18 and IL-6 ^34–36^ and cleave E-cadherin thereby weakening cell-cell interactions ^37^. Both meprins target cell junction proteins, thereby potentially enhancing leukocyte infiltration, degrade multiple ECM proteins (e.g. collagen IV, laminin-1, nidogen-1 and fibronectin) that stabilize the fibrous cap ^31,38^, and degrade lysyl oxidases that are critical for ECM assembly and maintenance ^39^. All of these processes are strongly associated with plaque development and progression thereby providing mechanistic links between elevated meprin protein and activity, increased inflammation, oxidant generation, loss of cell adhesion and communication, apoptosis, and weakening of the ECM within plaques. Meprin 1A also enhances tumor necrosis factor-α release by mast cells, and aggravates abdominal aortic aneurysms ^33^. Unlike MMPs and ADAMs, which can be activated by oxidants via the ‘cysteine switch’ ^40^ the pro-peptides of meprins are removed by trypsin-like proteases ^41^ including kallikreins (see below). Whilst these data are consistent with deleterious effects of elevated meprin levels and activity, these species also cleave human pro-collagen 1 at both N- and C-terminal sites (unlike bone morphogenic protein-1 which is C-propeptide specific) thereby facilitating collagen fibril formation ^42^.

Enhanced levels of caspases 1, 3, 6, 7 and 8 were detected in the cluster 2/3 datasets, with caspase 7 detected exclusively in these clusters. These enzymes play key roles in apoptosis (where caspase 8 is an initiator, and caspases 3, 6 and 7 are executioner enzymes), pyroptosis (with caspase 1 as an initiator) and necroptosis (caspase 8) ^43^. This family also convert pro-IL-1β ^44^ to active IL-1β, an important inflammatory mediator ^45^. IL-1β is the target of canakinumab, an antibody used in the CANTOS trial to demonstrate that dampening of inflammation has positive effects against cardiovascular disease ^46^. The over-abundance of meprins and caspases provides a link to enhanced IL-1β activation and canakinumab activity.

Cathepsins B, D, K and S have been implicated in atherosclerosis, particularly when present extracellularly ^47,48^. Cathepsin K and S deficiency decreases atherosclerosis in apoE^-/-^ mice, and these proteases have been identified by immunohistochemistry in human plaques ^49^. Elevated cathepsin K protein, as well as cathepsin B and S activity have been detected in symptomatic when compared to asymptomatic plaques ^48^. Enhanced cathepsin K levels have been detected in plasma after major adverse cardiovascular events, and to be an independent predictor of these ^50^. This cathepsin cleaves and activate pro-MMP-9 ^51^, linking cathepsins with MMP activation. Cathepsin D regulates apoptosis, and can modify APOB of LDL in a manner that promotes foam cell formation. Extracellular cathepsin B also degrades ECM proteins, and can enhance plaque vulnerability ^47^. Cathepsins L and S are potent collagenases and elastases ^52^, as is neurophil elastase (ELNE), with each of these over-abundant in cluster 2/3 plaques, consistent with greater ECM degradation. Furthermore, some of the peptide sequences present in the TopFIND database are consistent with elevated activities of these enzymes.

Some MMPs have been associated with plaque protein degradation ^28^. MMPs 3, 7, 11, 12 were over-expressed in clusters 2 and 3 (soft and mixed), whereas MMPs 8, 13 and 14 were primarily associated with cluster 1 (hard). Interestingly, MMP9, a major neutrophil-derived enzyme previously associated with plaque instability and rupture ^6,53–55^ was not detected as differentially expressed. The role of MMP3 appears to be complex and situation/species dependent, with enhanced levels associated with instability in humans ^56^ but stability in apoE^-/-^ mice ^57^. MMP12 has been reported to support plaque expansion and destabilization in mice ^57^. MMP7, when measured in plasma, has been reported to have neutral effects in mice ^57^ but to be associated with carotid stenosis in humans and to predict adverse outcomes ^58,59^. MMP13 inhibition enhances collagen deposition in mice, consistent with a role for this enzyme in ECM degradation ^60^. However, these data are inconsistent with the detected over-representation of MMP13 in the cluster 1 plaques, potentially indicating species dependent effects. MMP14 has been reported to be present at high levels in human plaques and to activate MMP2, amongst others ^61^. These divergent data may arise from the greater complexity of human plaques compared to murine disease, but also differences in samples (tissue versus plasma), detection methods (e.g. antibodies versus proteomics), and measurement of enzyme activity versus total protein. Thus, protein levels may not reflect activity in plaques, and one of the strengths of the current study is the combination of protein measurements ^16^ and peptides arising from enzymatic activity.

Several serine proteases were also elevated in clusters 2/3, including chymase (CMA1), a mast cell protein with ECM degrading activity ^62^. This enzyme generates vasoactive peptides, including (in place of angiotensin-converting enzyme) the conversion of angiotensin I to vasoactive angiotensin II ^62^. Enhanced chymase activity has been reported in internal thoracic arteries of people with hypocholesterolemia ^63^ and aneurysms ^64^. Serine protease HTRA2, was also detected at elevated levels. This mitochondrial and endoplasmic reticulum enzyme has been associated with caspase-dependent apoptosis and Parkinson’s disease, where it regulates (α-synuclein-mediated) mitochondrial oxidant production ^65^. This protein may therefore act in concert with caspases to enhance apoptosis.

Kallikrein-related peptidase 4 (KLK4) and plasmin/plasminogen (PLG/PLMN) were detected in all plaques, and not associated with particular clusters. KLK4 is a powerful activator of MEP1B ^66^ and may therefore contribute to enhanced MEP1B activity is soft plaques where this is elevated. Kallikrein enzymes are activated by a complex cascade involving multiple family members, and proteases of the thrombostasis axis including thrombin and plasmin ^67^. KLK5 was strongly associated with clusters 1 (hard) and 3 (mixed), and may contribute to ECM remodeling and wound repair ^68^. This enzyme cleaves E-cadherin, β-catenin, fibronectin and α5β1 integrin, in bronchial epithelial cells, and promotes epithelial-to-mesenchymal transition mechanisms and cell migration via the mitogen-activated protein kinase signaling pathway ^68^. KLK5 has also been associated with shedding of dipeptidyl peptidase-4 (DPP4 or CD26) from immune cells (particularly CD4+ T cells) into plasma, a process associated with the development of type 2 diabetes mellitus ^69^. Thrombin (THRB) was also overexpressed in cluster 1 plaques. This species has key pro- and anti-coagulant activities ^70^ and its enhanced levels may signify a greater importance of the latter activity in these plaques.

As many of the identified N-terminal peptides were absent from the topFIND database, Enrichment/Depletion sequence logo analysis was performed to uncover novel cleavage specificity patterns. A notable enrichment of Leu and Ser was detected (Figure 5C) at, and adjacent to, the P1’ position ^71^ with these residues commonly detected as the N-terminal residue in peptides over abundant in cluster 2 (Figure 5C, lower part). Val and Pro residues were also over abundant near the P1’ site, and particularly for cellular and core matrisome proteins (Figure 5C, left and middle panels). In contrast, for the matrisome-associated proteins, Ser residues were more enriched at P1’ for all plaque clusters (Figure 5C, right hand panels). The preference for Leu at P1’ is consistent with a role for MMPs (a common sequence specificity motif for this family and particularly MMPs 1, 2, 3, 7, 8, 12 and 14 ^72,73^) with several of these (MMPs 3, 7, 12) over-expressed in cluster 2, and MMPs 8 and 14 in cluster 1. The occurrence of Leu at P2’, and Pro at P3’ or P4’, is also consistent with MMP activity ^72,73^. The preference for Ser at P1’ is consistent with cleavage by Factor IX or thrombin isoforms ^74^ or ADAM family species (e.g. ADAM9). The mild preference for Ala at P1’ is consistent with Factor X/Xa cleavage ^75^ an enzyme over-represented in the cluster 1 (hard) plaques.

Analysis of the sites of cleavage on proteins yielding a large numbers of peptides (e.g. collagens 6A1, 6A2, 6A3 [COL6A1, COL6A2, COL6A3], complement component C3 [C3], kininogen 1 [KDG1], lumican [LUM], apolipoprotein B100 [APOB], and fibronectin [FN1]), indicates that there are significant differences in the peptides generated from specific proteins between plaque types, indicating that cleavage specificity is dependent on the plaque cluster (Supplementary Figure 1, where the cleavage sites for different plaque types are presented and color coded, and when determined, the protease responsible). These differences are exemplified by the data for COL6A3 and FN1, where similar numbers of peptides are detected for cluster 1 and clusters 2/3 but with different cleavage sites; C3 and KNG1, where many more cleavages are detected for cluster 1 and clusters 1/3; LUM, which is only detected as cleaved in clusters 2/3; and APOB, where many more peptides are detected for clusters 2/3, than cluster 1. The peptides abundant in, or exclusive to, cluster 2 or clusters 2/3, might be potential biomarkers of these plaque types and instability. Future studies will examine whether these species efflux into the circulation and urine.

## STRENGTHS AND LIMITATIONS

A major strength of this study is the breadth and depth of the N-terminal proteomic analyses. Of major importance is the strong correlation observed between the proteomes of the different clusters identified, and their gross morphological status (hard versus mixed and soft), ultrasound categorization (echolucent/echogenic) and presence of hemorrhage/ulceration. This N-terminal proteomic approach allows the identification of target proteins, specific proteases and the fragments that arise from enzymatic activity. Both known and novel proteases have been detected at both the protein, and activity level. The large numbers of novel neo-N-terminal species indicates the presence and action of novel proteases, and/or additional sequence specificities and protein targets to those established in the literature. The detection of fragments generated by particular proteases indicative of activity, rather than protein levels, is a key strength. This is of particular importance as many proteases are secreted in *inactive* forms, which may not be distinguished from active forms by gene analysis or antibody approaches.

This study has a number of weaknesses. One of these is the modest sample numbers, although these were sufficient for cluster determination and the detection of significant differences using multidimensional scaling and hierarchical clustering analyses. These data, therefore, provide a strong ‘proof-of-concept’, but do not allow analysis of other factors including age, sex, smoking and other factors (e.g. diabetes) that may influence plaque protein degradation. Only data from symptomatic, and not asymptomatic carotid stenosis, are included, as asymptomatic stenosis is not operated on in Denmark. Follow-up data from the subjects examined would also be of value, but is not available.

Further weakneses include a lack of plaque histology, as this is incompatible with the protein extraction protocol employed. Future studies should endeavor to resolve this issue, perhaps via prior plaque dissection. This study also lacks data from control healthy tissue, and information from early plaques, though such samples are difficult to obtain. Data from ‘healthy’ tissue would also allow discrimination between fragments from normal physiology and disease.

## CONCLUSIONS

This study provides valuable insights into the pathogenesis of atherosclerosis, focusing on the critical role of proteolysis in plaque destabilization. A novel N-terminal proteomics approach has been employed to identify proteases present in atherosclerotic plaques and the peptides that arise from their activity. Considerable numbers of novel peptides that cannot be assigned to known proteases have also been identified in these plaques, indicating that additional novel proteases are present. These data shed light on molecular events that may contribute to plaque destabilization, as many of the proteases and peptides are overabundant in plaques with soft and mixed macroscopic morphologies. These distinct plaque profiles highlight the heterogeneity of plaques, and a potential rupture-prone phenotype. Identifying key proteases and peptide fragments (and their protein substrates) unveils potential therapeutic targets for plaque stabilization. These data may also pave the way for an assessment of the propensity of plaques to undergo rupture, as the peptides may efflux from the plaques into plasma and urine. Together, these data emphasize the value of N-terminal proteomics in unravelling the proteolytic landscape of atherosclerotic plaques, and provides a foundation for further research aimed at combating atherosclerotic vascular disease.

## Nonstandard Abbreviations and Acronyms

ADAM: A Disintegrin And Metalloproteinase
APOB: apolipoprotein B-100
CEA: carotid endarterectomy
CVD: cardiovascular disease
DDA-PASEF: data-dependent acquisition with parallel accumulation-serial fragmentation
ECM: extracellular matrix
FDR: false discovery rate
GO: gene ontology
IL-1β: interleukin 1β
LC-MS: liquid chromatography-mass spectrometry
LC-MS/MS: liquid chromatography-mass spectrometry-mass spectrometry
LDL: low-density lipoproteins
MMPs: matrix metalloproteinases
MS: mass spectrometry
SPEED: Sample Preparation by Easy Extraction and Digestion protocol
TIMPs: tissue inhibitors of matrix metalloproteinases
US: ultrasound

## Acknowledgements

The authors thank the surgical teams of the Rigshopitalet, Capital Region, Copenhagen, for the collection of the endarterectomy samples.

## Sources of Funding

This work was funded by grants from Novo Nordisk Foundation grants NNF20SA0064214 (M.J. Davies) and and the Novo Nordisk Foundation-University of Copenhagen BRIDGE scheme (to L.G. Lorentzen).

## Disclosures

Prof. Davies reports declares consultancy contracts with Novo Nordisk A/S, and is a founder and shareholder in Seleno Therapeutics plc. These funders had no role in the design of the study; in the collection, analyses or interpretation of data; in the writing of the manuscript, or in the decision to publish the results. All other authors declare no conflicts of interest.

## Author Contributions

Drs Lorentzen and Davies have full access to all of the data in the study and take responsibility for the integrity of the data and the accuracy of the data analysis.

*Concept and design:* Lorentzen, Davies

*Acquisition, analysis or interpretation of data:* Lorentzen, Yeung, Eldrup, Eiberg, Davies

*Drafting of the manuscript:* Lorentzen, Davies

*Critical revision of the manuscript for intellectual content:* Lorentzen, Yeung, Eldrup, Eiberg, Davies

*Statistical analysis:* Lorentzen, Yeung

*Obtained funding:* Davies, Lorentzen

*Supervision:* Eldrup, Eiberg, Davies

## Supplemental Materials

Online Data Supplemental Table 1 and Supplemental Figures 1-2.

## Notes

### Competing Interest Statement

The authors have declared no competing interest.

